# Autophagy induction mitigates FUS aggregate formation and early synaptic dysfunction at the NMJ in the FUS-ALS model

**DOI:** 10.64898/2026.02.19.706635

**Authors:** Tulika Malik, Sam Jones, Olivia Ma, Sahana Mohan, R Michael Burger, Daniel T Babcock

**Affiliations:** Department of Biological Sciences, Lehigh University, Bethlehem, PA

## Abstract

Mutations in Fused in Sarcoma (FUS), a RNA binding protein, cause Amyotrophic Lateral Sclerosis (ALS). ALS is an aggressive neurodegenerative disease resulting in motor neuron degeneration. Defects in synaptic integrity precede neuronal loss in ALS, but the mechanisms responsible for these early synaptic defects are unclear. To investigate early synaptic defects associated with ALS, we expressed an ALS-linked variant of human FUS in adult motor neurons and assessed synaptic pathology at the neuromuscular junction (NMJ). Here we highlight the accumulation of FUS–positive aggregates at synaptic terminals and subsequent reduction in microtubule stability. We show that inducing autophagy via expression of Rab1 or Fragile-X Mental Retardation Protein 1 (FMR1), or treatment with Rapamycin reduces aggregate formation and restores synaptic structure and function. These findings reveal the utility of inducing autophagy to address early synaptic dysfunction in an ALS model and demonstrate a potential therapeutic target to preventing later stages of disease progression.

## Main

ALS is a fatal neurodegenerative disease characterized by the progressive loss of motor neurons and subsequent muscle atrophy^1^. As with other neurodegenerative diseases, disruption of synaptic integrity is an early hallmark of ALS that occurs long before neuronal loss^2^. However, the mechanisms underlying early synaptic defects that precede motor neuron loss are not well understood. We recently characterized early synaptic defects in a Drosophila model of ALS by expressing an ALS-linked FUS^3^. Over 50 mutations in *FUS* have been linked to ALS most of which function in a dominant fashion^4^. One of the most aggressive forms of ALS is linked to the FUS point mutation (FUS^R521C^), which is positioned at the C-terminal nuclear localization signal^5^.

ALS-linked FUS^R521C^ leads to motor neuron loss, functional defects, and cytoplasmic mislocalization in various models^6 7 8^. Recently, it was established that early synaptic defects at the NMJ precede this FUS-linked motor neuron loss, with disruption of the microtubule-associated protein 1B (MAP1B) among the earliest impaired synaptic markers^3^. MAP1B plays an important role in axon transport, which is known to be disrupted in TDP-43 models of ALS^9^. MAP1B has also been identified as a target of TDP-43, revealing disruption of microtubule dynamics in ALS pathology^10^. ALS-linked mutations in FUS lead to mislocalization of FUS from the nucleus to the cytoplasm, but it is unclear to what extent this mislocalization is responsible for the early synaptic defects that precede motor neuron loss.

One common link between FUS and other ALS models is autophagy disruption^11 12 13^. Both *FUS*^P525L^ and *FUS*^R522G^ mutations directly impair autophagy induction via LC3II levels in neuronal cell lines and primary cortical neurons^11^. Reduced autophagy seen with expression of ALS-linked FUS is also associated with accumulation of FUS-positive cytoplasmic stress granules in motor neurons, with Rapamycin treatment reducing stress granule formation^14^ ^15^.

Here we examine the mechanism underlying MAP1B defects and FUS pathology at the NMJ upon expression of FUS^R521C^. We uncover the accumulation of large FUS-positive aggregates at synaptic terminals that precedes microtubule impairment. Induction of autophagy via rapamycin treatment reduces synaptic aggregate volume and maintains microtubule stability. Additionally, we demonstrate that co-expression of Rab1 or Fragile-X Mental Retardation Protein 1 (FMR1) along with FUS^R521C^ restores microtubule defects and reduces synaptic aggregate volume. Furthermore, we report that inducing autophagy via expression of Rab1 or FMR1, or treatment with rapamycin, rescues defects in synaptic transmission at the pre and post-synapse. Collectively, our findings uncover the mechanism underlying early synaptic dysfunction and restoration of FUS-induced synaptic defects is autophagy dependent in our ALS model.

## Results

### ALS-linked FUS^R521C^ forms synaptic aggregates and disrupts cytoskeleton stability

To investigate synaptic pathology in adult Drosophila motor neurons upon expression of FUS, we expressed either wild-type or ALS-linked version of human FUS (FUS^WT^ or FUS^R521C^) in motor neurons innervating the Ventral Abdominal Muscles (VAMs). The synaptic architecture of these synapses resembles that of mammalian NMJs^16^. We recently demonstrated that ALS-linked FUS leads to early synaptic defects at these NMJs that precede motor neuron loss^3^.

To determine the mechanism underlying this synaptic dysfunction, we first assessed FUS localization of HA-tagged FUS^WT^ or FUS^R521C^ at synaptic terminals. We found that FUS^WT^ largely co-localizes with Futsch, a MAP1B homolog, at both Day 5 and Day 21 (Fig.1a & c). However, we observed accumulation of FUS-positive aggregates at the NMJ upon expression FUS^R521C^ (Fig.1b & d). These aggregates are present early at Day 5, with the average number and volume of aggregates holding steady at Day 21 (Fig.1e & f). Mislocalization of aggregate-prone FUS from the nucleus into the cytosol is an important hallmark of ALS^7^ . Our finding that this aggregate formation extends to the synaptic terminal suggests that these aggregates could underlie the early synaptic defects in our ALS model.

Our previous work identified disruption of cytoskeleton integrity at the NMJ as an early hallmark of synaptic dysfunction in this FUS ALS model, as indicated by a reduction in Futsch staining followed by the emergence of breakpoints throughout synaptic terminals^3^. To compare the onset of FUS aggregate formation and disruption of cytoskeleton integrity, we also measured Futsch breakpoints at synaptic terminals at Day 5 and Day 21 with the expression of FUS^WT^ and FUS^R521C^. At Day 5, there is a significant increase in the number of Futsch breakpoints at synaptic terminals with expression of FUS^R521C^, which progressively increases with age as seen at Day 21 (Fig. 1g-m). Together, these results demonstrate that FUS-positive aggregates at synaptic terminals accumulate alongside early defects to cytoskeleton integrity.

**Figure 1:**
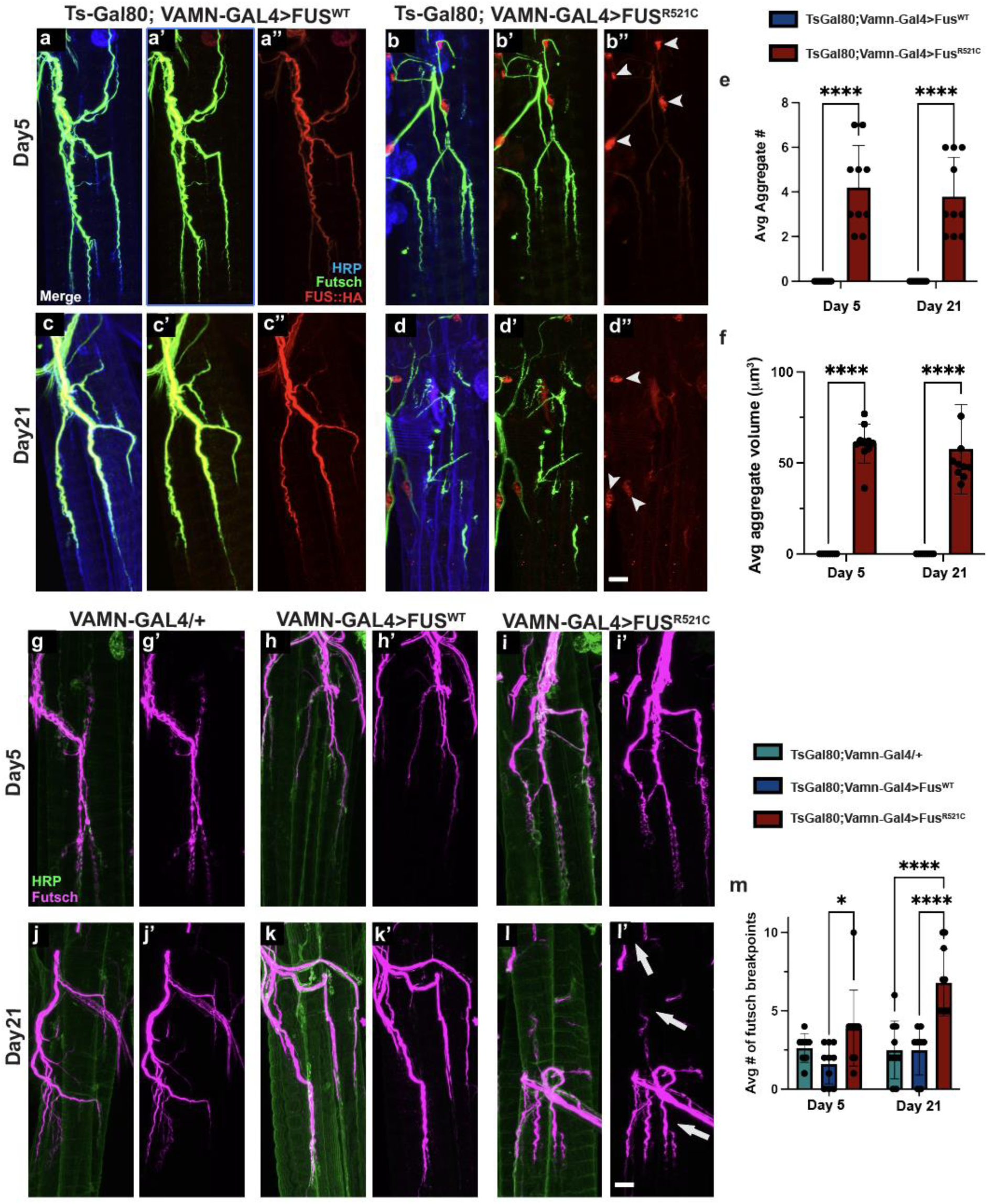
FUS^R521C^ forms large synaptic aggregates and disrupts cytoskeleton integrity. (a-d’’)Synaptic distribution of the microtubule-associated protein Futsch (green) and FUS (red) at Day 5 (a-b’’) Day 21(c-d’’) in controls or flies expressing FUS^WT^ or FUS^R521C^. Neuronal membranes are labeled with HRP (blue). Arrowheads indicate FUS-positive aggregates. (e-f) Average number (e) and volume (f) of FUS-positive-aggregates per NMJ. (g-l’) Synaptic distribution of Futsch (magenta) along with HRP (green) at the NMJ at Day 5 (g-i’) and Day21 (j-l’) for Vamn-gal4/+ controls, as well as flies expressing either FUS^WT^ or FUS^R521C^. Futsch breakpoints are shown with arrows. (m) Average number of Futsch Breakpoints per sample. Error bars represent the standard error of the mean (sem). *p<0.05 , ****p<0.0001, using one-way ANOVA with Tukey’s Post hoc test for multiple comparisons. Scale Bars in d’’ and l’ are 10μm.

To further investigate the disruption of cytoskeleton integrity in this ALS model, we tested whether the breakpoints seen in Futsch staining indicate severing of Tubulin or reduced microtubule stability. To assess severing of Tubulin, we stained motor neurons expressing either FUS^WT^ or FUS^R521C^ for alpha-Tubulin at Day 5 and Day 21. We did not observe any significant increase in alpha-Tubulin breakpoints in flies expressing either FUSWT or FUS^R521C^ compared to controls at either early or late timepoints (Fig. 2a-g). This suggests that the breakpoints seen in MAP1B staining do not indicate a severing of microtubules.

**Figure 2:**
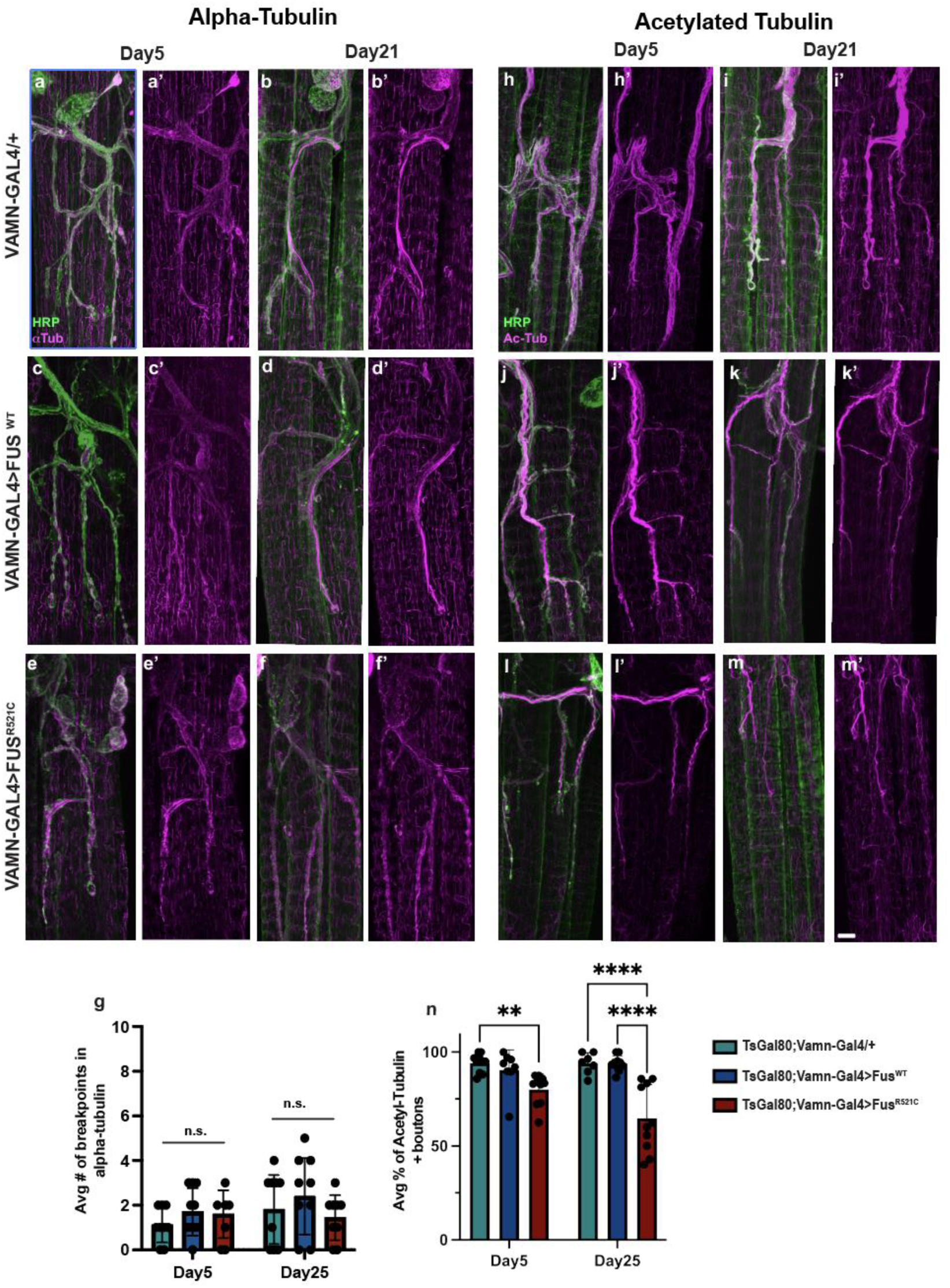
Expression of FUS^R521C^ disrupts microtubule stability. (a-g) Distribution of Alpha-Tubulin (magenta) and HRP (green) at Day 5 and Day21 for Vamn-gal4/+ controls (a-b’), or flies expressing FUS^WT^ (c-d’), or FUS^R521C^(e-f’). (g)Average number of breakpoints in Alpha-Tubulin per sample. (h-n) Distribution of Acetylated-Tubulin (magenta) and HRP(green) at Day 5 and Day 21for Vamn-gal4/+ controls (h-i’), or flies expressing FUS^WT^ (j-k’) or FUS^R521C^(l-m’). (n)Percentage of Acetylated-Tubulin positive boutons per sample. Error bars represent the standard error of the mean (sem). **p<0.01, ****p<0.0001, n.s.= not significant, using one-way ANOVA with Tukey’s Post hoc test for multiple comparisons. Scale Bar in m’ is 10μm.

Since microtubules do not appear to be severed, we next assessed microtubule stability at motor neuron terminals by measuring the synaptic distribution of acetylated-Tubulin, a well-known marker of stable microtubules^17^. We found that flies expressing FUS^R521C^ exhibited a significant decrease in the percentage of synaptic boutons that stained positive for acetylated-Tubulin at Day 5, which progressively worsened by Day 21. The reduction in acetylated-Tubulin staining was specific for ALS-linked FUS^R521C^ , since this phenomenon was not observed in flies expressing FUS^WT^ or in controls (Fig. 2h-n). Together, these results demonstrate that the early synaptic defects observed upon expression of FUS^R521C^ are due to reduced microtubule stability through disruption of MAP1B.

### Rapamycin treatment reduces FUS-positive synaptic aggregates and maintains microtubule stability

To investigate the mechanisms responsible for these early synaptic defects, we tested whether treatment with rapamycin can mitigate this aggregate formation and disruption of microtubule stability. rapamycin is an inhibitor of Mammalian Target of Rapamycin (mTOR) that is well known to induce autophagic flux in neurons and enhance degradation of aggregate-prone proteins^18^. Autophagy disruption has been shown in multiple ALS models such as TDP-43, FUS, and SOD1^19^. Additionally, rapamycin’s therapeutic effects have been well established in TDP-43 models of ALS^20^ .

To assess whether rapamycin treatment rescues synaptic defects upon expression of FUS^R521C^, we measured aggregate volume and MAP1B breakpoints in flies expressing either FUS^WT^ or FUS^R521C^ raised on either standard media or media supplemented with 300μM Rapamycin. We found a significant reduction in average aggregate volume at both Day 5 and Day 21 in Rapamycin treated FUS^R521C^ - expressing flies compared to those without Rapamycin treatment (Fig. 3a-e). We also observed a significant reduction in MAP1B breakpoints with rapamycin treatment when compared to controls (Fig.3f). Finally, we assessed cytoskeleton stability by measuring the percentage of acetylated-tubulin positive boutons with and without rapamycin treatment. We observed that rapamycin treatment significantly restores the percentage of acetylated-tubulin positive boutons in FUS^R521C^ expressing flies back to normal at both Day 5 and Day 21 (Fig. 3g-k). Additionally, we measured p62 accumulation at the NMJ with expression of Fus^WT^ or Fus^R521C^. We observed that Fus^R521C^ expression results in significantly higher p62 intensity when compared to controls (Extended Data Fig. 1a-b’). Furthermore, when treated with rapamycin, the p62 intensity is rescued back to normal (Extended Data Fig. 1c-d). This accumulation of p62 suggests an increase of polyubiquitinated proteins upon expression of Fus^R521C^, which is rescued with autophagy induction in our FUS-ALS model. Together, these results indicate that treatment with rapamycin can mitigate the formation of early synaptic aggregates, prevent destabilization of microtubules, and reduce P62 accumulation.

**Figure 3:**
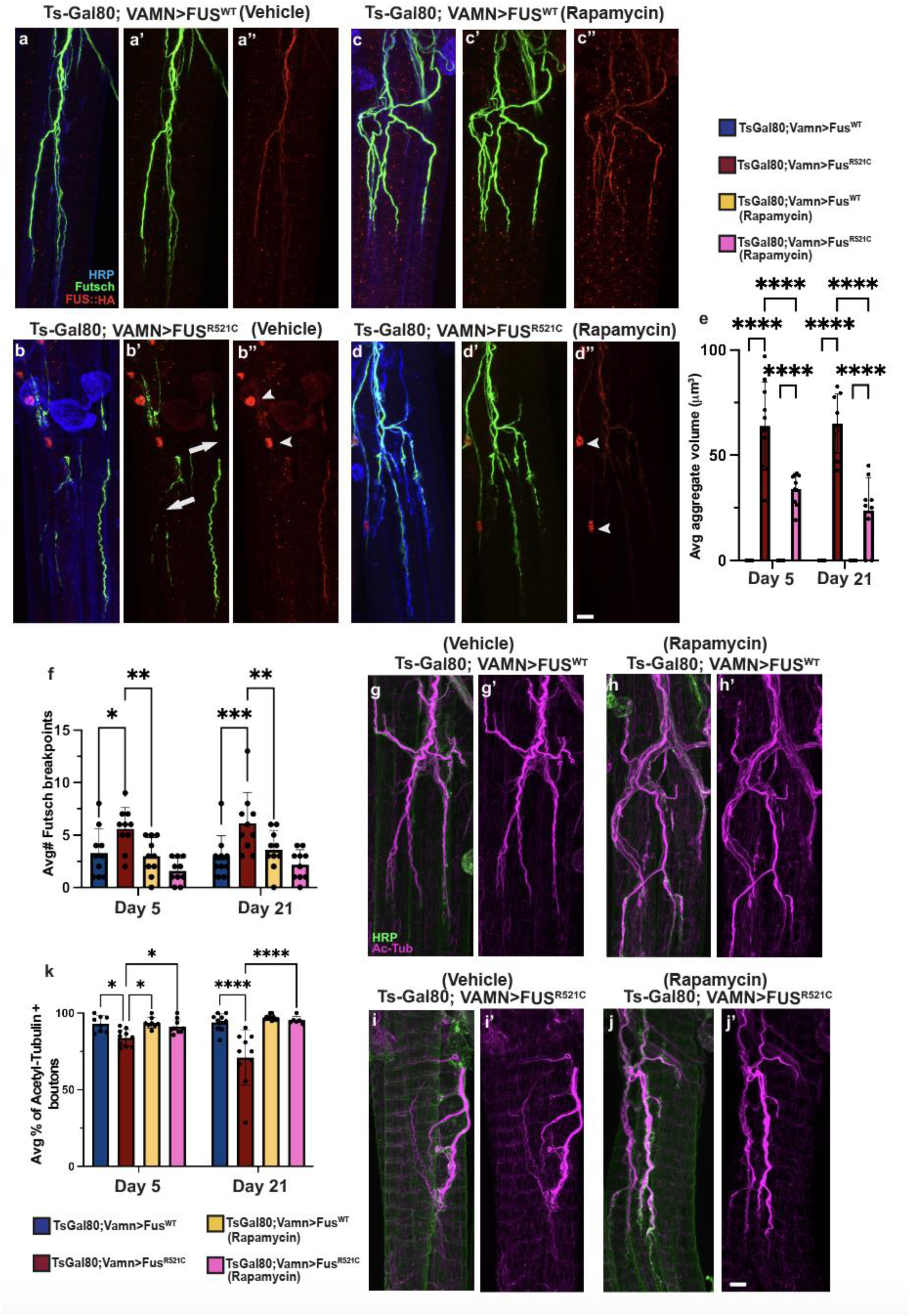
Rapamycin treatment reduces FUS-positive aggregate volume and maintains cytoskeleton stability. (a-f) Synaptic distribution of the microtubule-associated protein Futsch (green), FUS (red), and HRP (blue) at Day 21 when raised on standard food (a-b’’) or on food supplemented with rapamycin (c-d’’) in flies expressing FUS^WT^ or FUS^R521C^. Arrowheads indicate FUS-positive aggregates, and arrows indicate breakpoints in Futsch staining. (e) Average aggregate volume for vehicle and rapamycin treated groups at Day 5 and Day 21. (f) Average number of Futsch breakpoints for vehicle and rapamycin treated groups at Day5 and Day21. (g-k) Distribution of Acetylated-Tubulin(magenta) and HRP(green) at Day 21 for flies raised on standard food with vehicle only (g-h’) or on rapamycin-supplemented food(i-j’) in flies expressing FUS^WT^ or FUS^R521C^. (k) Average percentage of Acetylated-Tubulin positive boutons at Day 5 and Day 21 for all groups. Error bars represent the standard error of the mean (sem). *p<0.05 , **p<0.01, ***p<0.001, ****p<0.0001 using one-way ANOVA with Tukey’s Post hoc test for multiple comparisons. Scale Bar in d’’ and j’ are 10μm.

### FMRP or Rab1 expression in motor neurons reduces synaptic aggregates and maintains microtubule stability in the FUS ALS model

Fragile X Mental Retardation protein (FMRP) is associated with ALS pathology^21^, with FMRP levels decreasing upon expression of mutant FUS or TDP-43^21 22^. Like rapamycin, FMRP is an mTOR inhibitor and mediates autophagy similarly in Fragile X syndrome models^23^. Since FMRP and rapamycin both work by blocking mTOR, we investigated whether we can replicate the pharmacological rapamycin-mediated rescue by overexpressing FMRP while expressing FUS^WT^ or FUS^R521C^ . To test if FMRP expression can mitigate synaptic aggregates and microtubule impairment, we expressed FMRP along with FUS in adult motor neurons. We observed that FMRP expression prevents the formation of FUS^R521C^-induced synaptic aggregates (Fig. 4a-e). Additionally, we found that MAP1B breakpoints are similar to those in controls upon co-expression of FUS^R521C^ and FMRP at both Day5 and Day21(Fig. 4a-f). Lastly, we revealed that FMR1 overexpression maintains cytoskeleton stability by restoring the percentage of acetylated Tubulin-positive boutons (Fig. 4g-k). These results indicate FMRP expression maintains cytoskeleton stability and prevents synaptic aggregate formation at the NMJ in this FUS model of ALS.

**Figure 4:**
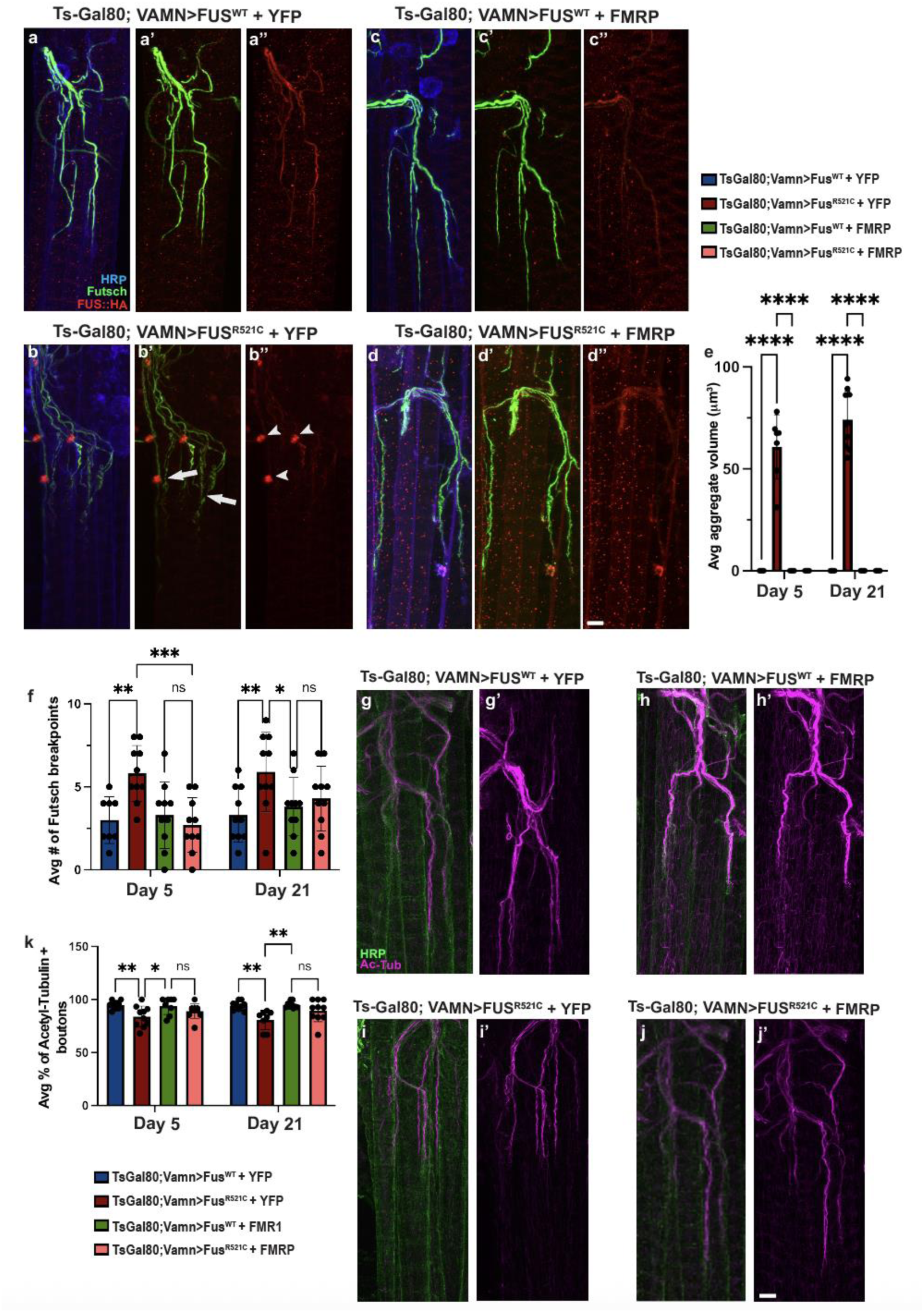
FMRP expression prevents FUS-positive aggregate formation and maintains cytoskeleton stability in FUS^R521C^. (a-d’’) Synaptic distribution of the microtubule-associated protein Futsch (green), FUS(red), and HRP(blue) together at Day 21 while co-expressing EYFP (a-b’’) or FMR1(FMRP)(c-d’’) along with FUS^WT^ or FUS^R521C^. Arrowheads indicate FUS-positive aggregates, and arrows indicate breakpoints in Futsch staining. (e) Average aggregate volume with and without the overexpression of FMRP at Day 5 and Day 21. (f)Average number of Futsch breakpoints with and without the expression of FMRP at Day 5 and Day 21. (g-j’) Distribution of Acetylated-Tubulin (Magenta) and HRP (Green) at Day 21 while co-expressing EYFP (g-h’) or FMR1(FMRP) (i-j’)along with FUS^WT^ or FUS^R521C^. (k)Average percentage of Acetylated-Tubulin positive boutons at Day 5 and Day 21 for all groups. Error bars represent the standard error of the mean (sem). *p<0.05 , **p<0.01, ***p<0.001, ****p<0.0001, n.s.=not significant, using one-way ANOVA with Tukey’s Post hoc test for multiple comparisons. Scale Bars in d’’ and j’are 10μm.

Our results demonstrate that both pharmacological and genetic inhibition of mTOR signaling rescue early synaptic defects in our ALS model. In addition to the induction of autophagy, mTOR signaling is also involved in translational repression^24^. To test whether mTOR-independent induction of autophagy can similarly rescue early synaptic defects, we next assessed microtubule stability and synaptic aggregate formation upon expression of Rab1. Rab1 is one of several Rab GTPases that are specifically involved in autophagosome formation^25^. Previous studies have shown that autophagosome formation is disrupted in ALS-models, and increased expression of Rab1 restores these defects^11^. To test if Rab 1 expression can mitigate synaptic aggregates and microtubule impairment in our ALS model, we expressed Rab1 along with FUS specifically in adult motor neurons. We measured aggregate volume and MAP1B breakpoints in flies expressing either FUS^WT^ or FUS^R521C^ while expressing Rab1. Similar to our results with Rapamycin treatment and FMRP expression, we found a reduction in average aggregate volume at both Day5 and Day 21 in FUS^R521C^ overexpressing Rab1 (Fig. 5a-e). We also observed a significant reduction in MAP1B breakpoints in FUS^R521C^ overexpressing Rab1 when compared to controls (Fig. 5a-f). This MAP1B breakpoint reduction is seen at Day5 and remains consistent at Day21. Finally, we assessed cytoskeleton stability by measuring the percentage of acetylated-tubulin positive boutons in FUS^WT^ or FUS^R521C^ while expressing Rab1. We found that Rab1 expression prevented the reduction of acetylated tubulin-positive boutons in adult flies expressing FUS^R521C^ at both Day 5 and Day 21. Together, these results suggest that inducing autophagy in an mTOR-independent manner via expression of Rab1 reduces aggregate volume and limits cytoskeleton defects at the NMJ in our FUS model of ALS.

**Figure 5:**
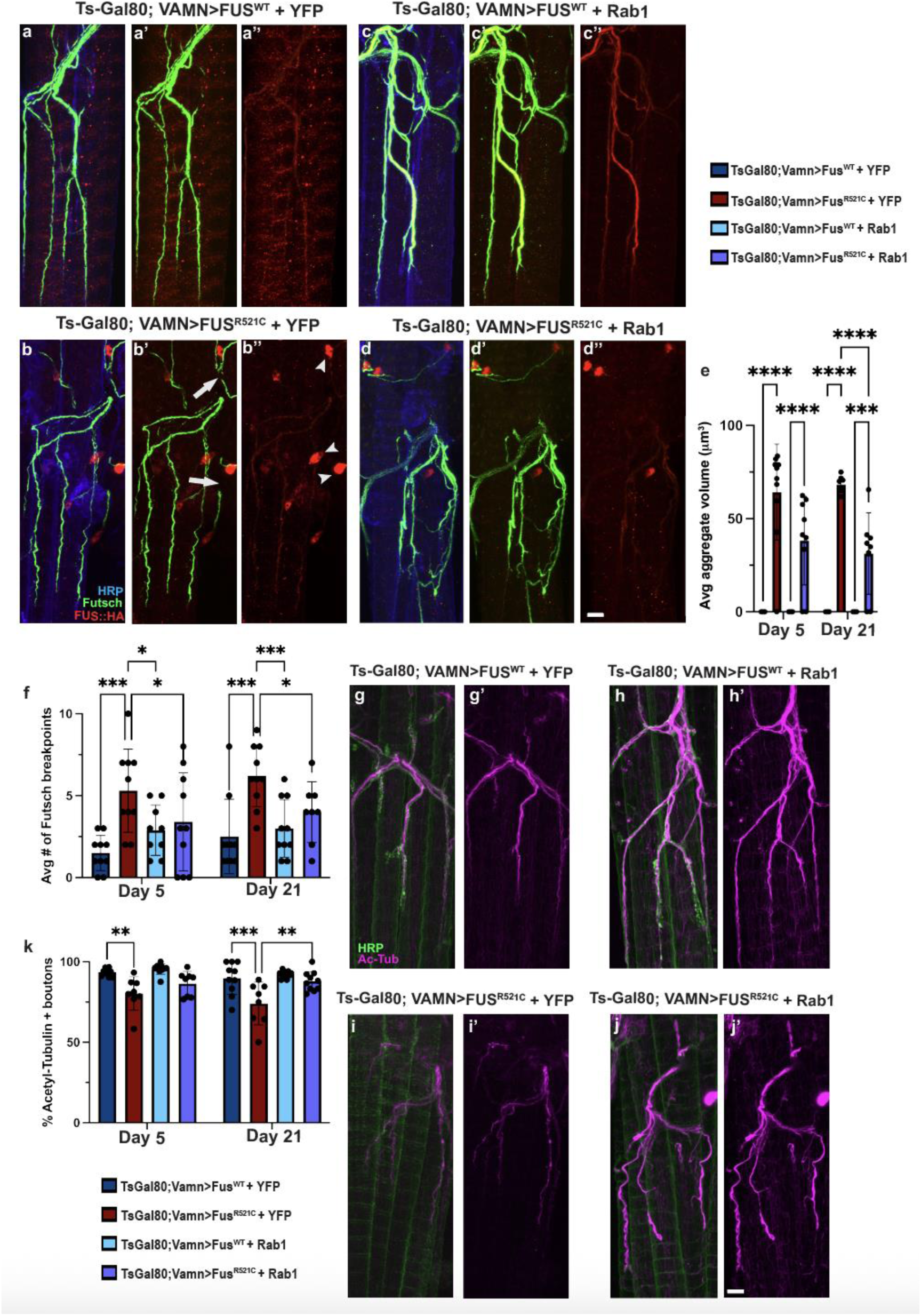
Rab1 expression reduces FUS-positive aggregate volume and maintains cytoskeleton stability in FUS^R521C^. (a-d’’) Synaptic distribution of the microtubule-associated protein Futsch (green), FUS (red), and HRP (blue) at Day 21 while co-expressing EYFP (a-b’’) or Rab1(c-d’’) along with FUS^WT^ or FUS^R521C^. Arrowheads indicate FUS-positive aggregates, and arrows indicate breakpoints in Futsch staining. (e) Average aggregate volume with and without the overexpression of Rab1 at Day 5 and Day 21. (f)Average number of Futsch breakpoints with and without expression of Rab1 at Day5 and Day21. (g-j’) Distribution of Acetylated-Tubulin (magenta) and HRP (green) at Day 21 while co-expressing EYFP (g-h’) or Rab1(i-j’) with FUS^WT^ or FUS^R521C^. (k) Average percentage of Acetylated-Tubulin positive boutons at Day 5 and Day 21 for all groups. Error bars represent the standard error of the mean (sem). *p<0.05 , **p<0.01 ***p<0.001 ****p<0.0001, using one-way ANOVA with Tukey’s Post hoc test for multiple comparisons. Scale Bars in d’’ and j’are 10μm.

### Autophagy induction rescues synaptic and motor function in the FUS^R521C^ model

We next assessed whether conditions that rescue synaptic structure and disrupt aggregate formation similarly maintain synaptic function in our ALS model. We previously described early defects in abdominal curling behavior in flies expressing FUS^R521C^ in motor neurons innervating the Ventral Abdominal Muscles^3^. Using optogenetics, we stimulated the Ventral Abdominal Motor Neurons (VAMNs) and recorded the average percentage of curling behavior in flies expressing FUS^WT^ or FUS^R521C^ at Day 5 and Day 21. Similar to our previous study, here we confirm that FUS^R521C^ causes a significant decline in curling compared to controls and FUS^WT^ at Day 5 and Day 21(Fig. 6a). Importantly, in the current study we further show that treatment with rapamycin restores abdominal curling behavior to normal levels in files expressing FUS^R521C^ at Day 5 and Day 21(Fig. 6a). This indicates that rapamycin treatment maintains both structural and function integrity of synapses in this FUS-linked ALS model.

**Figure 6:**
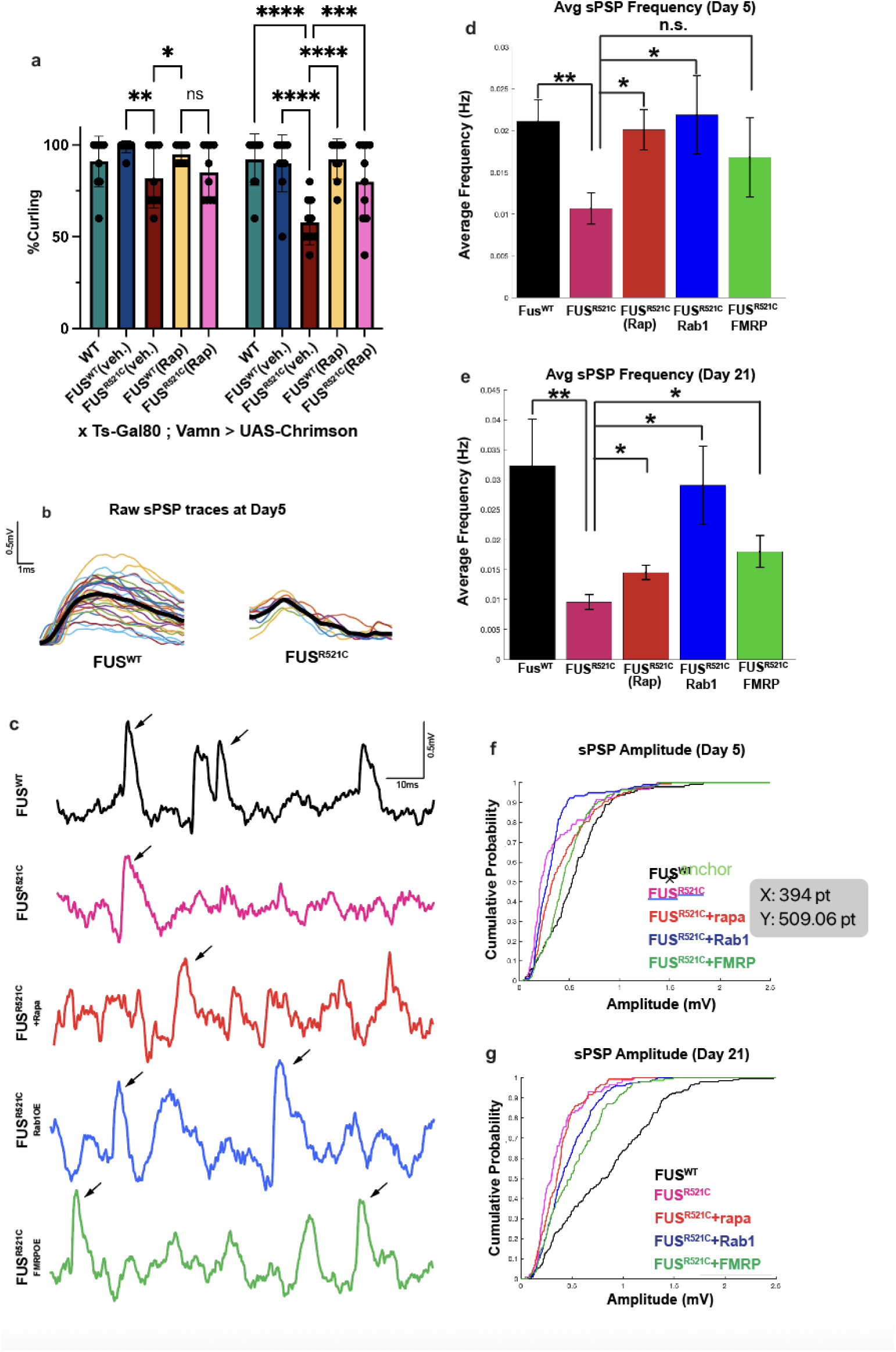
Autophagy induction rescues synaptic and motor function in the FUS^R521C^ model. (a)Measurement of the percentage of stimulations that induced abdominal curling at Day 5, and 21. Genotypes include motor neuron expression of UAS-Chrimson along with wildtype FUS (Blue), FUS^R521C^ (red), wildtype FUS with rapamycin treatment (yellow), FUS^R521C^ with rapamycin treatment (pink) or Luciferase (green) as a control for transgene expression. Each dot represents the average response rate for an individual fly. *p<0.05 , **p<0.01 ***p<0.001 ****p<0.0001, using one-way ANOVA with Tukey’s Post hoc test for multiple comparisons. (b) Single traces illustrate sPSP events at Day5 for flies expressing either FUS^WT^ or FUS^R521C^. The average trace is represented by the *black line*. (c) sPSP traces at Day 21 for wildtype FUS (black), FUS^R521C^(magenta), FUS^R521C^with rapamycin treatment (red), FUS^R521C^with Rab1 overexpression (blue), and FUS^R521C^with FMRP overexpression (green). (d-e) Average sPSP frequency at Day5 and Day21 for all genotypes (n=10), *p<0.05 using two-sample t-test on MATLAB. (f-g) Cumulative probability of sPSP indicates significant reductions in sPSP amplitude in the FUS^R521C^ model at Day5 and Day21(n=10), *p<0.05 using a two-sample Kolmogorov-Smirnov test. Cumulative plots show significant restoration of sPSP amplitude after rapamycin treatment or FMRP OE models at Day 5. While at Day 21 a significant restoration of sPSP amplitude after RAB1 OE or FMRP OE models, and a partial restoration for rapamycin treatment is observed.

To further investigate the synaptic transmission defects induced by FUS^R521C^, we recorded postsynaptic potentials at the NMJs of both Day 5 and Day 21 for all treatment groups. To first confirm that the observed events were synaptic, recordings at the adult control NMJ showed that they were abolished by DNQX application and recovered after a 10-min washout (Extended Data Fig. 2a–c′). We then recorded spontaneous postsynaptic potentials (sPSPs) using sharp intracellular electrodes in current-clamp mode in flies expressing FUS^WT^ or FUS^R521C^. We observed a significant decline in sPSP frequency at both Day5 and Day21 for files expressing FUS^R521C^ compared to FUS^WT^(Fig. 6c-e). Interestingly, treatment with rapamycin or either expression of FMRP or Rab1 restored the average frequency in flies expressing FUS^R521C^ at Day 5 and Day21 (Fig. 6d-e).

We also observed that when FUS^R521C^ is expressed, the cumulative amplitude probability curve shifts significantly to the left at both Day 5 and Day 21, indicating significantly smaller amplitudes in this FUS model of ALS (Fig 6.f-g). At both timepoints we observed a significant rescue in the population amplitudes as the curves shifted toward the control values, indicating that sPSP amplitudes are also partially or fully rescued for all three rescue models (Fig 6.f-g). The reductions in both sPSP frequency and amplitude that we observed are consistent with both pre- and postsynaptic defects (respectively) in synaptic transmission when FUS^R521C^ is expressed at the motor neuron. We further demonstrated complete or partial rescue of these effects with either rapamycin treatment or with overexpression of Rab1 or FMRP.

### Rescue of early synaptic defects by rapamycin is autophagy dependent

Finally, we tested the hypothesis that the rescue mediated by rapamycin treatment is dependent on its promotion of autophagy within motor neurons. To examine this, we blocked autophagy in motor neurons by knocking down ATG5, an essential protein involved in autophagosome formation^26^, while treating them with rapamycin. We assessed whether rapamycin treatment is still able to rescue FUS-induced synaptic defects, while ATG5 knockdown. We analyzed both aggregate volume and microtubule disruption at Day 5 and Day 21. Aggregate volume did not decrease upon knockdown of ATG5 even with rapamycin treatment (Fig. 7a-d), consistent with an autophagy-dependent mechanism of rapamycin rescue. Similarly, we found that rapamycin treatment reduced the number of FUS^R521C^ - induced microtubule breakpoints. However, ATG5 knockdown prevented the rapamycin-induced rescue of Futsch breakpoints at Day5 and Day21 (Fig. 7a-e). Together, these results show that blocking autophagy prevents rapamycin from exerting its neuroprotective effects in this FUS-ALS model.

**Figure 7:**
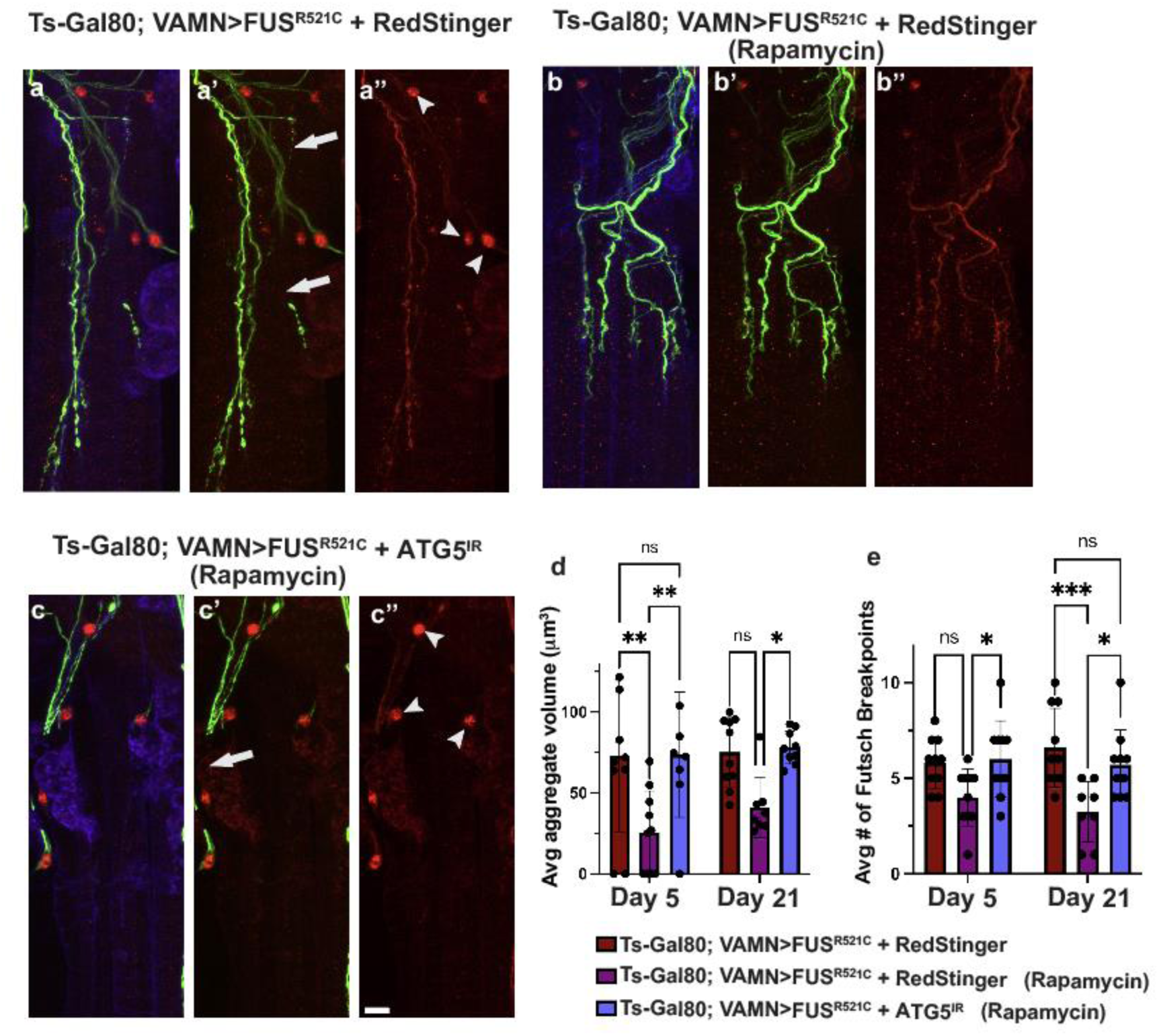
Rapamycin induced rescue of early synaptic defects requires autophagy. (a-c’’) Synaptic distribution of the microtubule-associated protein Futsch (green), FUS (red), and HRP (blue) at Day 21 while co-expressing redstinger as a control (a-a’’) or ATG5^IR^ on rapamycin supplemented food (b-b’’) or redstinger on rapamycin supplemented food (c-c’’) along with FUS^R521C^. (d) Average aggregate volume with and without ATG5 KD at Day 5 and Day 21. (e)Average number of Futsch breakpoints with and without ATG5 KD at Day 5 and Day 21. *p<0.05, **p<0.01, ***p<0.001, n.s.=not significant, using one-way ANOVA with Tukey’s Post hoc test for multiple comparisons. Scale Bar in c’’is 10μm.

## Discussion

Advances in ALS research have demonstrated a strong link between autophagy disruption and ALS progression. Our current study demonstrates that autophagy induction specifically within presynaptic motor neurons exerts a neuroprotective effect in our FUS-ALS model. Autophagy is essential for clearing excess, mislocalized, or misfolded proteins from the cell. Multiple ALS models have shown autophagy disruption including the FUS-ALS model^13 14^. Autophagy disruption in FUS-ALS models can further lead to downstream effects such as mitochondrial dysfunction, DNA damage, and defects in RNA metabolism^19^. Here, we show that autophagy induction maintains synaptic integrity in this FUS-ALS model.

Synaptic defects are among the earliest known hallmarks of ALS, with disruptions to synaptic structure and function preceding motor neuron loss by a wide margin^27^. Characterizing the earliest synaptic defects in this ALS model and determining how synaptic integrity can be restored is a crucial step for developing neuroprotective strategies. A major objective of this study was to determine whether restoration of early synaptic defects prevents or delays ALS progression.

ALS-linked mutations in FUS are associated with the accumulation of mislocalized FUS within the cytoplasm, resulting in motor neuron defects^28 6 7^. However, the mechanism by which FUS mislocalization results in early synaptic defects is unclear. In this study, we showed that expressing human mutant FUS within motor neurons leads to the formation of FUS-positive aggregates at the synaptic terminal. Additionally, we revealed early cytoskeleton defects that align well with other TDP-43 ALS model studies^10^.

Further, we revealed that rapamycin treatment in the FUS-ALS model reduces synaptic aggregate formation, cytoskeleton defects, and P62 accumulation at the NMJ. Rapamycin has also been shown to be protective in TDP-43 and FUS ALS models^20^ , suggesting that the effects are likely widespread. Finally, we demonstrated that overexpression of the mTOR inhibitor, FMRP, which is reduced in multiple ALS models and is known to regulate MAP1B ^23 22 31^, similarly rescues early synaptic pathology. Since rapamycin and FMRP are both translation repressors, we decided to test Rab1, a mTOR-independent autophagy induction pathway. Overexpression Rab1 in motor neurons also reduced synaptic aggregate volume and cytoskeleton defects. Lastly, we showed that blocking autophagy through ATG5 knockdown prevents rapamycin-induced protective effects in the FUS-ALS model, demonstrating that the pharmacological and genetic rescue of early synaptic defects work via autophagy induction.

Our work reveals that the FUS-ALS model leads to disruption of both the structural and functional integrity of synapses. We find that behavioral defects occur alongside physiological defects in our ALS model. Our findings indicate physiological impairments at both pre- and postsynaptic structures. Although previous studies have shown presynaptic defects in Drosophila larvae using both FUS and C9orf72 ALS models^28 32^, the postsynaptic defects uncovered here provide crucial insights into the early synaptic pathology in this ALS model. Induction of autophagy similarly rescued synaptic function, indicating that autophagy plays an important role in mitigating ALS pathology by preserving NMJ structure and function.

Uncovering the mechanisms underlying these earliest stages of synaptic pathology is crucial for understanding disease progression. Our work highlights the potential of autophagy induction as an important ALS target to mitigate this FUS-induced early pathology. This will serve an important role in formulating early and effective therapeutic strategies.

## Methods

### Drosophila Stocks and Husbandry

All fly stocks were maintained on standard Drosophila medium. To provide temporal control of Gal4-induced transgene expression, we used a temperature-sensitive Gal80 (*Tub-Gal80^TS1^*)^33^ . Gal4 activity was inactive at 18°C and Gal4 activity was activated at 29°C. As previously described^3^,crosses were made at 18°C and remained until eclosion, at which point adult progeny were collected and shifted to 29°C for aging.

The following fly stocks were obtained from Bloomington Drosophila Stock Center (BDSC): *24G08-Gal4* (#49316)^34^, which we refer to as V*AMN-Gal4*^35^, *Tubulin-Gal80^TS^*^1^ (#7018)^33^ *20xUAS-Chrimson::Venus* (#55135)^36^, *UAS-Luciferase* (#35788)^37^ , *UAS-EYFP* (#6659)^38^, *UAS-Rab1-YFP* (#24104)^39^, *UAS-FMR1-GFP* (#92833)^40^, and *UAS-ATG5^IR^* (#34899)^41^. UAS-FUS^WT^-HA and UAS-FUS^R521C^-HA were kindly provided by Udai Pandey^8^.

### Immunohistochemistry

Ventral Abdominal Muscles (VAMs) were dissected as previously described (Malik et al., 2025). Briefly, the abdomen was isolated and cut through the dorsal midline, followed by pinning the cuticle flat and removal of fat body tissue. Samples were fixed in 4% paraformaldehyde in PBS for 30 minutes, then washed four times with PBS at room temperature (RT). Next, tissues were incubated in Blocking Buffer (PBS with 0.1% normal goat serum and 0.2% Triton X-100) for at least 1 hr at 4°C .

Samples were incubated overnight at 4°C with the following primary antibodies: mouse anti-Futsch (22C10) (1:200) and mouse anti-alpha-Tubulin (4A1) (1:100) from the Developmental Studies Hybridoma Bank, rat anti-HA (3F10) (1:400) (Sigma-Aldrich), rabbit anti-p62 (AB178440) (1:200) (Abcam), and mouse anti-acetylated-Tubulin (6-11B-1)(1:250) (Millipore).

Samples were then washed four times in PBS with 0.3% Triton X-100 (PBS-T) for at least 5 minutes followed by incubation in secondary antibodies. Samples were incubated in the following secondary antibodies for 2 hours at RT in the dark: anti-HRP 488 (1:200), anti-HRP 647 (1:25), Goat anti-Mouse 488 (1:200), Goat anti-Mouse 568 (1:200) and Goat anti-Rat 568 (1:200). Finally, samples were washed four times for 5 mins each with PBS-T and mounted on a glass slide with Vectashield (Vector Laboratories).

### Confocal Microscopy

Images were acquired using a Zeiss LSM 880 confocal microscope. Adult VAM samples were imaged using a 63x oil objective with a scanning depth of 1.0μm per optical slice. Brightness and contrast for images were adjusted using ImageJ software (NIH)^42^ and Adobe Photoshop. All figures were generated in Adobe Illustrator (Adobe Creative Cloud 2025).

### Aggregate Volume extraction

syGlass software (syGlass Inc.) was used to analyze the volume of FUS-positive aggregates. Czi files from the confocal were uploaded. Using the ROI tool in syGlass, aggregates were filled, and mask data was created. Mask data of aggregates was used to extract aggregate volume.

### Rapamycin Treatment

For Rapamycin treatment flies were raised on standard food supplemented with 300 μM of Rapamycin (Fisher Scientific). Rapamycin was dissolved in ethanol and then mixed directly into the food.

### Optogenetics

Optogenetic stimulation of Ventral Abdominal Motor Neurons was performed as previously described^3^ .*20xUAS-Chrimson::Venus* was expressed in motor neurons using Vamn-Gal4 along with *Tubulin-Gal80^TS1^*. Flies also expressed either *UAS-FUS^WT^*, *UAS-FUS^R521C^*, or *UAS-Luciferase* as a control. Crosses were made on standard media at 18°C, and flies were collected after eclosion. Flies were raised at 29°C in the dark on standard media supplemented with 0.25 mM All trans-retinal (Spectrum Chemical). Flies were anesthetized on ice for 10 minutes and the thorax was glued to a glass slide (Elmer’s Clear Glue). Glued flies were allowed to acclimate in the dark for 20 minutes. Flies were stimulated for 10 seconds with 623nm light with an illuminator (Wayllshine), followed by 30 seconds of rest. This process was repeated for a total of 10 stimulations per fly, and the percentage of stimulations resulting in abdominal curling were measured. Optogenetic experiments were recorded using a 4K digital video camera with an infrared lens (Zohulu). These experiments were performed in the dark under a wide angle infrared illuminator (Lonnky, model L3030A).

### Electrophysiology

Adult flies were anesthetized on ice for 1 minute before dissection. Flies were pinned down in the sylgard dish, and dissections were performed at room temperature in 0 mM Ca^2+^ physiological saline HL3.1. Electrophysiology dissections were performed by first pinning the head and dissecting down the midline while keeping the thorax and abdomen undamaged^43^. Make sure to remove fat bodies and trachea before placing them in the recording chamber. Replace 0 mM saline solution with recording extracellular saline containing 1.5 mM Ca^2+^, HL3.1 saline solution for recording^44^. The extracellular solution contained in mM (70 NaCl, 5 KCl, 1.5 CaCl_2_, 4 MgCl_2_ , 10 NaHCO_3_, 5 Trehose, 115 Sucrose, 5 HEPES) with 309 mOsm. Sharp (15-20 MΩ) borosilicate glass intracellular electrodes were pulled using a pipette puller (Narishege, PC-10) and filled with 3M KCl. Recordings were conducted from A2 and A3 hemi-segments only. Recordings were conducted in cells with sustained baseline membrane voltage below -40mV. During recording epochs, slow time constant current injections were applied to hold membrane voltage between -45 mV to -50 mV. sPSP in current-clamp mode. Recordings in each fly did not exceed 30 mins. sPSPs were detected by thresholding the zeromeaned and lightly smoothed (5-8 pts) voltage signal. Threshold was a scalar (2-4) of the variance of the voltage signal. Smoothing and threshold parameters were individually chosen for each recording to optimize sPSP selection. sPSP frequency and amplitude were calculated from 10 sec recording epochs. Custom MATLAB code is available upon request.

## Acknowledgements

The authors would like to thank members of the Babcock and Burger Labs for their support and guidance. The authors would like to thank Dr. Brian McCabe and Dr. Samuel William Vernon for their guidance to optimize adult NMJ electrophysiology. We would like to thank Dr. Julie Haas for guidance and optimization with the electrophysiology analysis using MATLAB. We would also like to thank Dr. Udai Pandey for sharing the FUS transgenic lines.

---

**Extended Data Fig. 1:**
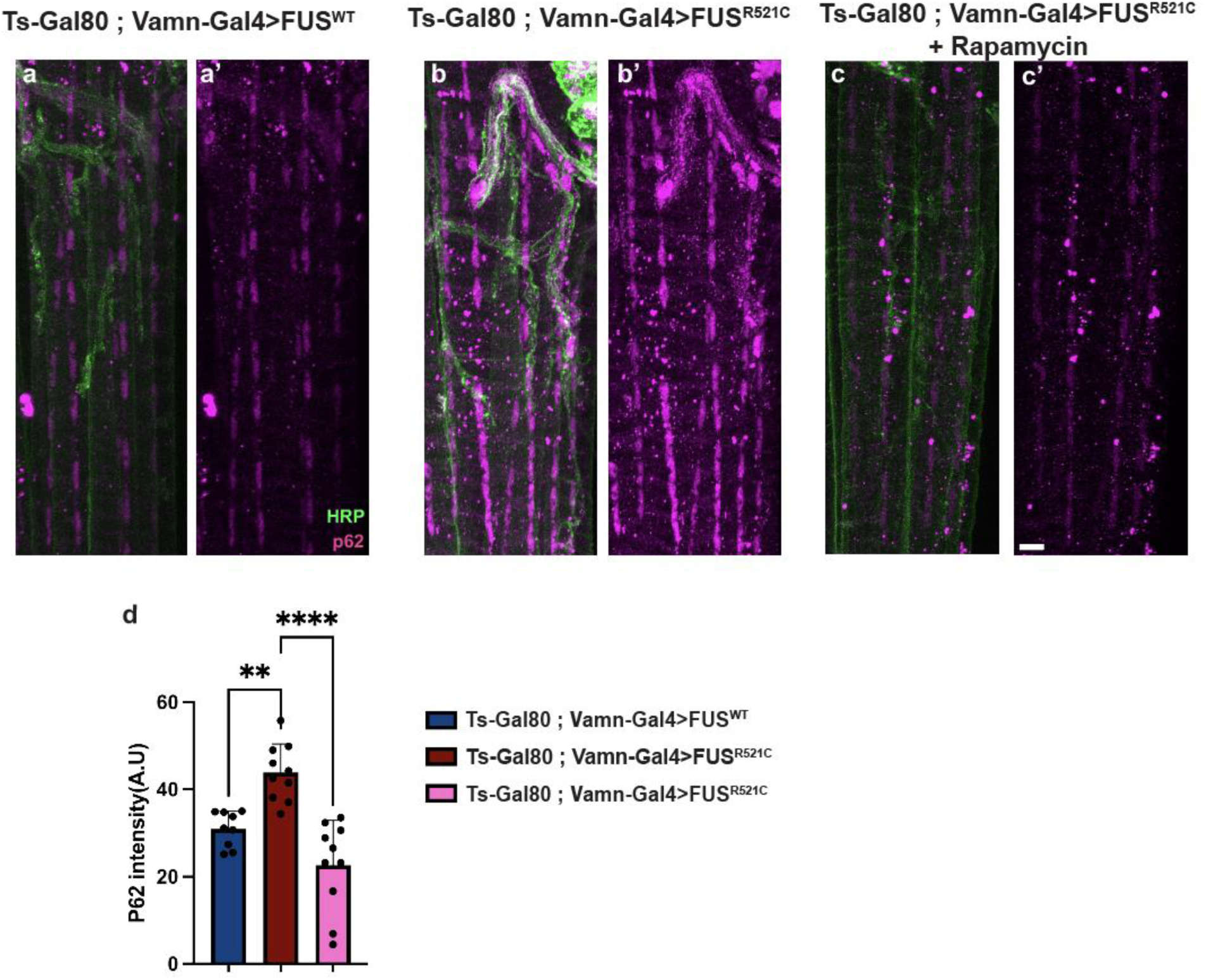
p62 accumulation at the NMJ in the FUS^R521C^ model is rescued with rapamycin treatment. (a-c’) P62 accumulation at the NMJ with the expression FUS^WT^ (a-a’), FUS^R521C^ (b-b’), and FUS^R521C^ on rapamycin-supplemented food (c-c’), p62 (magenta) and HRP (green). (d) p62 intensity for all three genotypes. **p<0.01 and ****p<0.0001 using one-way ANOVA with Tukey’s Post hoc test for multiple comparisons. Scale Bar in c’’ is 10μm.

**Extended Data Fig. 2:**
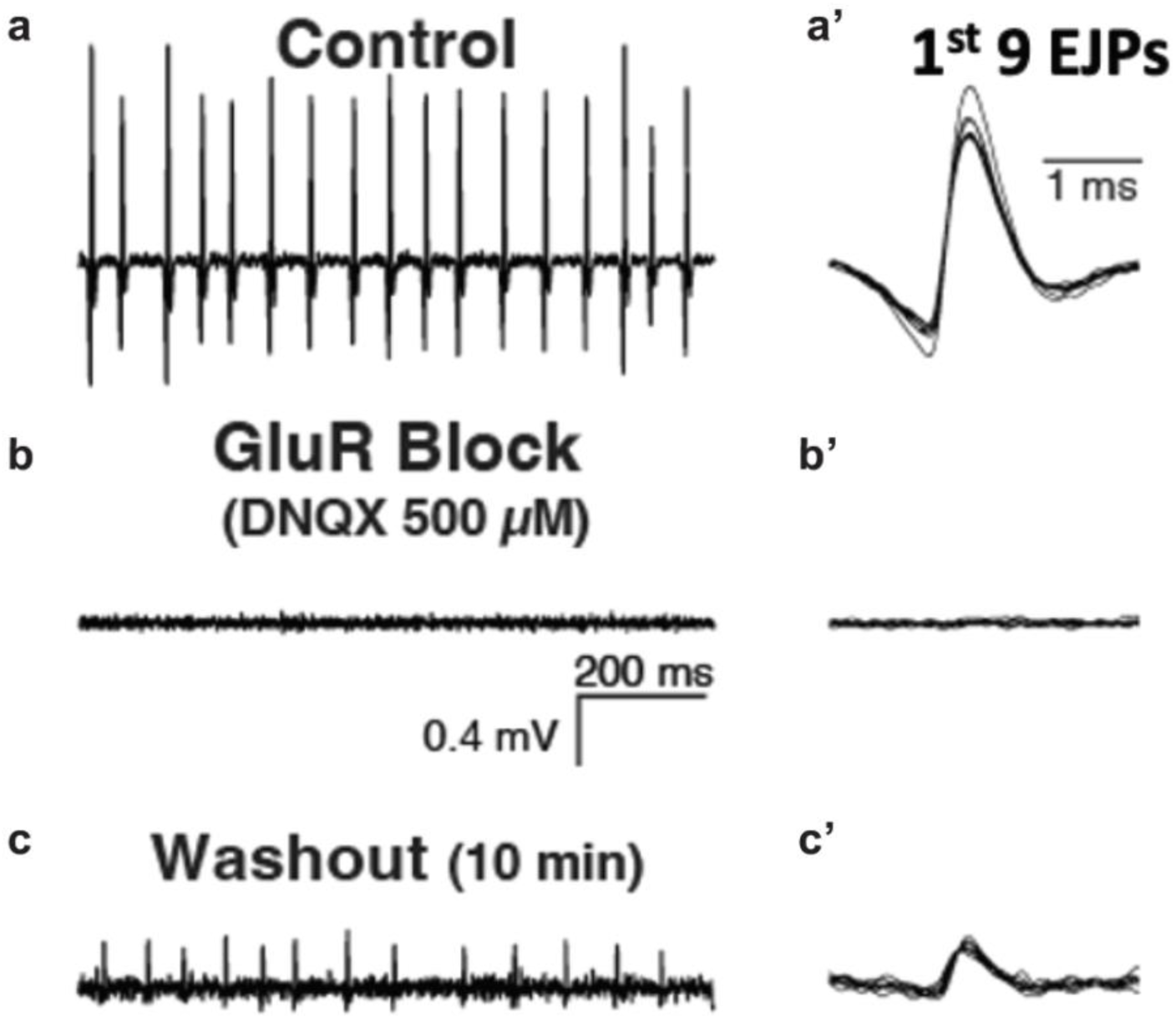
Blocking GluRs abolishes EJPs in control flies and recovered after washout. (a-c’) EJPs in control flies and the 1^st^ 9 EJPs before (a-a’), during DNQX application (b-b’) , and 10 mins after washout(c-c’).

